# Conservation Genomics of Urban Populations of Streamside Salamander (*Ambystoma barbouri*)

**DOI:** 10.1101/2021.11.04.467341

**Authors:** N. Wade Hubbs, Carla R. Hurt, John Niedzwiecki, Brian Leckie, David Withers

## Abstract

In Tennessee, populations of the state endangered Streamside Salamander (*Ambystoma barbouri*) are in decline as their distribution lies mostly within rapidly developing areas in the Nashville Basin. Information regarding the partitioning of genetic variation among populations of *A. barbouri,* and the taxonomic status of these populations relative to northern populations and their congener, the smallmouth salamander (*A. texanum*), have important implications for management and conservation of this species. Here we combined mitochondrial sequencing and genome-wide single nucleotide polymorphism (SNP) data generated using Genotyping-by-Sequencing (GBS) to investigate patterns of genetic variation within Tennessee populations of *A. barbouri*, to assess their relationship to populations in Kentucky, and to examine their phylogenetic relationship to the closely related *A. texanum*. Results from phylogenetic reconstructions reveal a complex history of Tennessee *A. barbouri* populations with regards to northern populations, unisexual *A. barbouri,* and *A. texanum*. Patterns of mitochondrial sequence variation suggest that *A. barbouri* may have originated within Tennessee and expanded north multiple times into Kentucky, Ohio, Indiana and West Virginia. Phylogenetic reconstructions based on genome-wide SNP data contradict results based on mitochondrial DNA and correspond to geographic and taxonomic boundaries. Variation in allele frequencies at SNP genotypes, as identified by multivariate analyses and Bayesian assignment tests, identified three evolutionary significant units (ESUs) for *A. barbouri* within the state of Tennessee. Collectively, these results emphasize the need for prioritizing conservation needs of Tennessee populations of *A. barbouri* to ensure the long-term persistence of this species.

## Introduction

Genetic variation, population structure and demographic history are increasingly recognized as important factors for the design of effective conservation strategies (Geist 2010). Streamside salamander (*Ambystoma barbouri,* Fig. 1) populations in Middle Tennessee are declining due to rapid urbanization in and around the Nashville Basin and as a result, were reclassified from "deemed in need of management" to state endangered by the Tennessee Wildlife Resources Agency (TWRA) (Withers et al. 2009, TWRA 2018, Anderson et al. 2014). Population fragmentation and loss of genetic variation that inevitably accompany loss of habitat threaten the long-term adaptive potential and persistence of *A. barbouri,* but can be mitigated by management actions aimed at maintaining genetic diversity. Information regarding *A. barbouri’s* taxonomy, population structure and patterns of genetic variation are currently needed to prioritize conservation needs and to efficiently allocate management resources. Genomic tools are increasingly being used to improve recovery and management planning in at-risk salamander species by informing taxonomic relationships, demographic histories, and biologically meaningful units of conservation that will preserve genetic diversity and the long-term adaptive potential of species.

**Figure 1.**
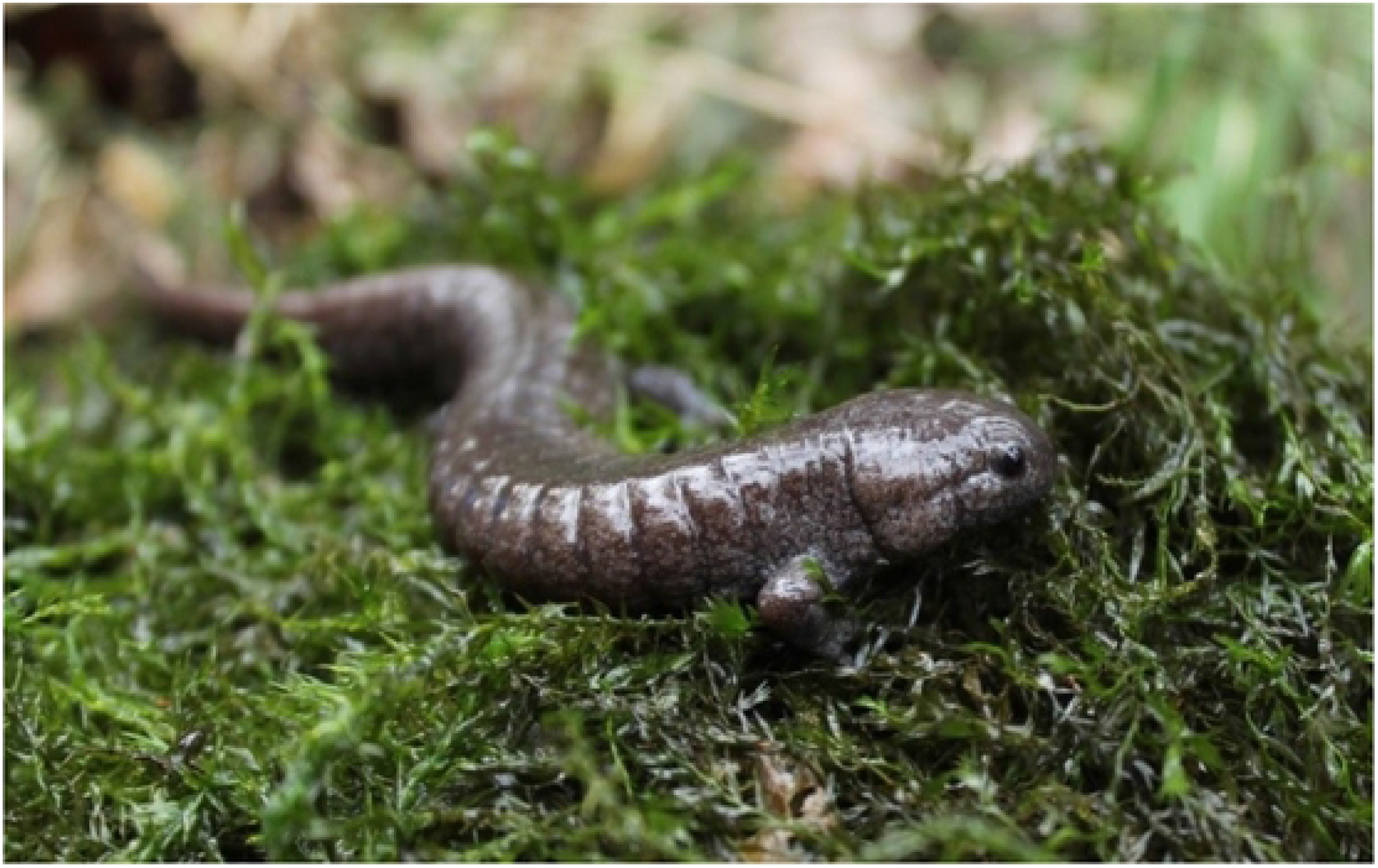
Photo image of Tennessee *Ambystoma barbouri*.

Taxonomic uncertainties surrounding the evolutionary relationship between *A. barbouri* and its sister-species *A. texanum* (smallmouthed salamander) are a concern for precisely identifying targets of conservation. These two species are nearly indistinguishable based on external morphological features alone and were previously considered to be conspecific. They can be differentiated using scanning electron microscopy on the number and shape of maxillary and premaxillary teeth, life-history traits, and choice of breeding environment (Niemiller et al. 2009, Kraus & Petranka 1989). In addition to their morphological ambiguity, phylogenetic reconstructions using mitochondrial and nuclear sequence data also produced conflicting results regarding the relationship between *A. texanum* and *A. barbouri* (Niedzwiecki 2005).

Reconstructions based on mitochondrial sequences show that *A. texanum* and *A. barbouri* are not reciprocally monophyletic; *A. texanum* is recovered as a clade nested within *A. barbouri* from Tennessee, suggesting that pond-breeding *A. texanum* were more recently derived from a stream-breeding *A. barbouri* ancestor from Central Tennessee. However, reconstructions based on two nuclear gene sequences resulted in reciprocally monophyletic clades for the two species, consistent with a history where *A. barbouri* and *A. texanum* are older independent lineages derived from a shared common ancestor. Additional sequence data are needed to resolve unanswered phylogenetic questions as these earlier reconstructions were based on a limited number of molecular markers (two nuclear and one mitochondrial gene) and representative outgroups.

Mitochondrial evidence also informs on the origins of unisexual Ambystomatid populations that are common in the Great Lakes region of North America. Unisexual Ambystomatids exhibit a unique reproductive mode whereby male sperm activates egg development, but contributes variable amounts of nuclear genetic material depending on compatibility with unrelated cytoplasmic DNA (termed kleptogenesis). While the nuclear genome unisexuals is a mixture of different Ambystomatid species, all known mitochondrial haplotypes, across their range, nest within *A. barbouri.* Mitochondrial haplotypes are most closely related to mitochondrial haplotypes found south of the Ohio River, and southwest of the Kentucky river (Bogart et al. 2007); however, the relationship of Tennessee *A. barbouri* to unisexual Ambystomatids has not been explored.

At a finer scale, uncertainties also exist regarding the relationship among isolated populations of *A. barbouri* in Tennessee and discontinuous populations in Kentucky, Indiana, Ohio, and West Virginia. Phylogenetic reconstructions of *A. barbouri* populations based on mitochondrial sequence data (913 bp) from the D-loop and an adjacent intergenic spacer suggest that Tennessee *A. barbouri* populations are both ecologically and genetically distinct from more northern populations (Eastman et al. 2009). At hatching, *A. barbouri* from Tennessee were smaller and less developed than individuals from Kentucky (Niedzwiecki 2005). Also, laboratory behavioral assays show that Tennessee *A. barbouri* were similar to western Kentucky individuals, but less active than individuals from northern populations (Niedzwiecki 2005). These differences were supported by genetic data; phylogenetic reconstructions recovered mitochondrial haplotypes from Tennessee as monophyletic and basal to haplotypes from Kentucky, Ohio, and West Virginia (Eastman et al. 2009). These results support an early divergence of Tennessee populations and raise questions regarding the geographic origin of *A. barbouri*. However, conclusions from this study were limited as these reconstructions are based on a single mitochondrial gene from only three individuals from a few populations within a kilometer of each other in Rutherford County.

Observations from field surveys suggest that *A. barbouri* populations in Tennessee are in decline, and the accompanying loss of genetic variation in small populations further threatens the long-term persistence of this species (Niemiller et al. 2006). Maintenance of genetic variation is a fundamental priority for conservation planning and requires information regarding the partitioning of genetic variation within and between isolated populations. Data on patterns of genetic variation in Tennessee populations of *A. barbouri* are limited to a handful of mitochondrial and nuclear sequences used for phylogenetic studies (Eastman et al. 2009; Niedzweicki 2005). Genome-wide surveys of genetic variation from fine-scale population sampling across the state are critical for assessing genetic variation within populations, establishing units of conservation and maintaining historical patterns of gene flow between populations to improve the outcome of recovery efforts (Shaffer et al. 1996).

Here we investigate patterns of genetic variation within and among populations of Tennessee *A. barbouri* as well as the taxonomic relationship between *A. barbouri* and *A. texanum* in order to prioritize conservation needs and inform management practices aimed at maximizing long-term persistence of this species. Mitochondrial sequence data and genome wide SNP genotypes were used to (1) investigate the phylogenetic relationship between *A. barbouri* and *A. texanum*, (2) review the taxonomic relationship of disjunct populations of *A. barbouri* in Tennessee relative to northern populations, (3) evaluate patterns of genetic differentiation between geographically isolated populations of *A. barbouri* within the state of Tennessee and (4) estimate within-population genetic variation and examine demographic history of Tennessee *A. barbouri*. Taxonomic relationships, units of conservation, and geographic partitioning of genetic variation are discussed in the context of establishing conservation priorities and designing effective management strategies for this species.

## Methods

### Tissue Collection and DNA Extraction

Tissue samples of *A. barbouri* in the form of adult tail clips, eggs, and whole larvae were obtained from field surveys or from collaborators (Fig. 2, Table 1). Historical and predicted sites of *A. barbouri* were surveyed from January 2018 to March 2018 and again from November 2018 to March 2019, coinciding with timing of oviposition as reported by Niemiller et al. (2009). At each site, *A. barbouri* adults and eggs were collected by turning over cover objects within and near seasonal streams. Additionally, pools were searched for free-swimming larvae; only one larva was sampled from any single pool to avoid sampling related individuals from the same clutch. A total of 235 individuals were included for genetic analysis. Samples included 225 individuals from 13 populations of *A. barbouri* in Tennessee spanning six counties in the Nashville Basin: Bedford County (B6), Davidson County (D3), Rutherford County (R1, R7, and R9), Sumner County (S2, S5, S7, S8), Wilson County (W1, W3, and W4), and Williamson County (Wil2). A total of five *A. texanum* individuals were sampled; these included four individuals from (Arnold Airforce Base, AAFB) and one from Craighead Co., Arkansas (AKT1). Outgroups included two *A. mabeei* (North Carolina), two *A. maculatum* (AAFB), and one *A. talpoideum* (AAFB).

**Figure 2.**
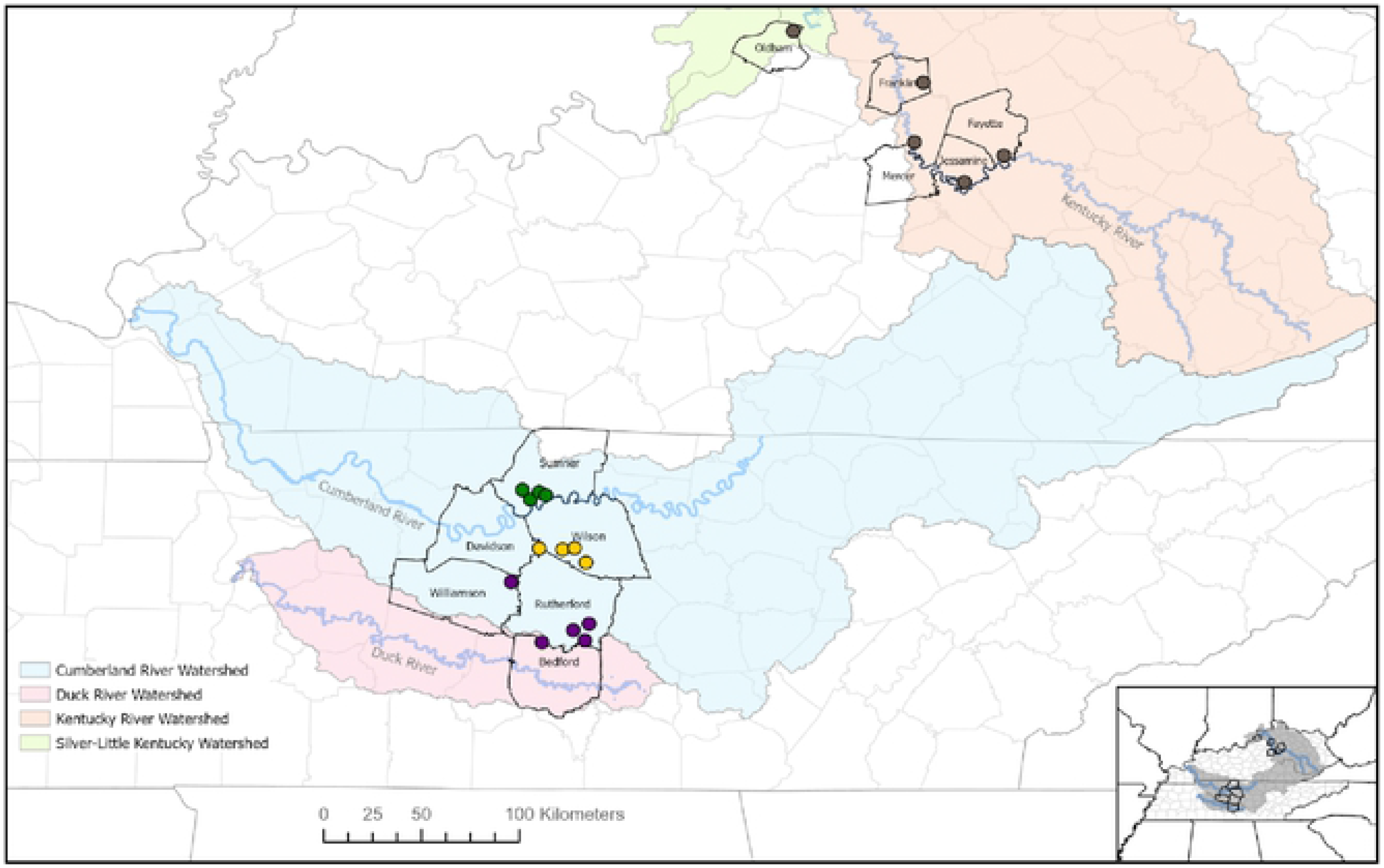
Map of all surveyed populations of *A. barbouri* in Tennessee and Kentucky. County lines are shown with black borders. Shaded regions indicate major watersheds as depicted in the legend. Tennessee populations are coloredby ESU assignment where green circles indicate northern ESU populations, yellow circles indicate central ESU populations, and purple circles indicate southern ESU populations.

**Table 1.** Population IDs, sample sizes for mtDNA/GBS analyses, and map coordinates for all populations. Population IDs for *A. barbouri* collections correspond to map locations in Figure 1. For outgroup samples, AAFB denotes Arnold Air Force Base and AKT1 denotes *A. texanum* from Arkansas. Population IDs by county are as follows: Bedford County (B6), Davidson County (D3), Rutherford County (R1, R7, and R9), Sumner County (S2, S5, S7, and S8), Wilson County (W1, W3, and W4), and Williamson County (Wil2).

| Population ID (Map ID) | Sample Size<br>mtDNA/GBS | Latitude | Longitude |
| --- | --- | --- | --- |
| <u>*A. barbouri* - Tennessee</u> |  |  |  |
| Sumner 2 (S2) | 3/-- | 36.336487 | -86.502042 |
| Sumner 5 (S5) | 3/11 | 36.329443 | -86.570217 |
| Sumner 7 (S7) | 5/37 | 36.363339 | -86.537930 |
| Sumner 8 (S8) | 6/22 | 36.367510 | -86.591100 |
| Davidson 3 (D3) | 3/29 | 36.100821 | -86.535713 |
| Wilson 1 (W1) | 5/15 | 36.040205 | -86.321052 |
| Wilson 3 (W3) | 4/12 | 36.093415 | -86.372530 |
| Wilson 4 (W4) | 3/6 | 36.093249 | -86.398001 |
| Williamson 2 (Wil2) | 3/5 | 35.950021 | -86.648606 |
| Rutherford 1 (R1) | 3/20 | 35.719718 | -86.353396 |
| Rutherford 7 (R7) | 12/28 | 35.672435 | -86.315247 |
| Rutherford 9 (R9) | 3/11 | 35.752811 | -86.306184 |
| Bedford 6 (B6) | 4/8 | 35.669700 | -86.528800 |
| <u>*A. barbouri* - Kentucky</u> |  |  |  |
| Kentucky FC (FC) | 3/3 | 37.772370 | -84.569400 |
| Kentucky (RR) | 5/4 | 37.897710 | -84.393000 |
| Kentucky SL (SL) | 4/3 | 38.490100 | -85.359400 |
| Kentucky SW (SW) | 4/5 | 38.251190 | -84.753400 |
| Kentucky JS (JS) | 2/2 | 37.960310 | -84.818100 |
| <u>Outgroups</u> |  |  |  |
| *A. mabeei* | 2/ -- | 33.936860 | -78.567865 |
| *A. maculatum* (AAFB) | 2/ -- | 35.392500 | -86.099722 |
| *A. talpoideum* (AAFB) | 1/1 | 35.392500 | -86.099722 |
| *A. texanum* (AAFB) | --/4 | 35.392500 | -86.099722 |
| *A. texanum* (AKT1) | 1/1 | 35.828812 - | -90.688313 |

DNA was extracted from tail clips, eggs, and larvae using the EZNA Tissue DNA Mini kit (OMEGA BIO-TEK) following the manufacturer’s protocol, except that DNA was eluted in water. Approximately 1200 bp of the mitochondrial D-loop was targeted for PCR amplification using primers developed by Shaffer and McKnight (1996; Table 2). The mitochondrial D-loop has been shown to be informative for evaluating population structure and species relationships within the genus *Ambystoma* (Bogart et al. 2007; Charney et al. 2014; Church et al.; 2003; Shaffer & McKnight 1996; Zamudio & Savage 2003). Conditions for polymerase chain reaction (PCR) were as follows: initial denaturation step of 2 minutes at 95°C, followed by 35 cycles of 15s at 95°C, 15s at 53°C, and 90s at 72°C. This program ended with a final extension of 10 min. at 72°C. Amplified PCR product was cleaned prior to cycle sequencing by exonuclease I/shrimp alkaline phosphatase (New England Biolabs) and used for bi-directional Sanger sequencing on an ABI 3730 automated sequencer (MCLAB). Sequence chromatograms were imported and visualized using SEQUENCHER 5.2 (Gene Codes Corporation). Sequences were aligned using ClustalW Multiple Alignment option (Thompson et al. 2003) as implemented in Bioedit (Hall 1999).

**Table 2.**
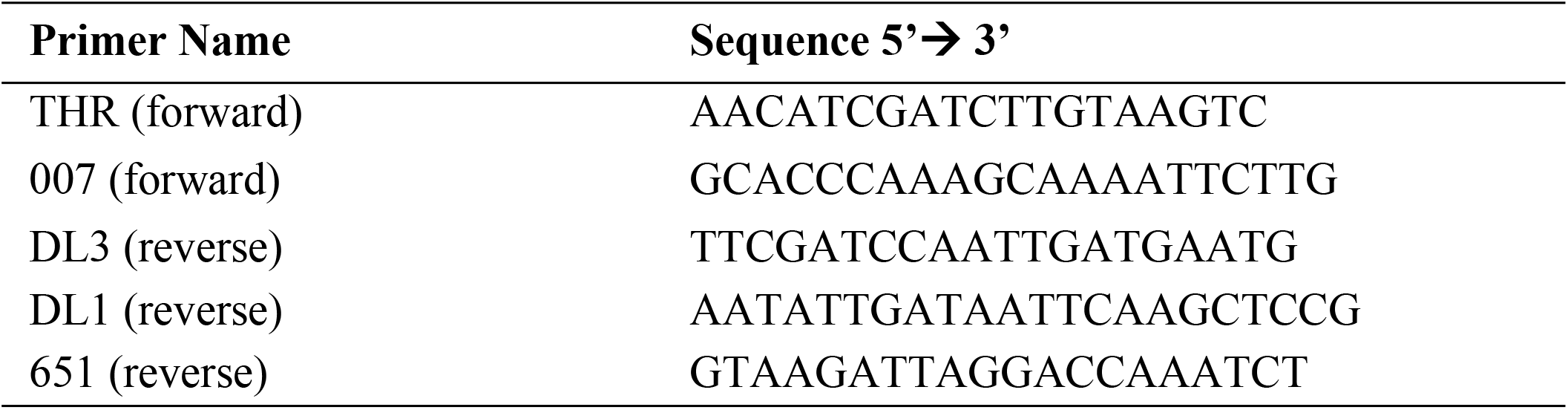
Primers used for PCR amplification of the mitochondrial D-loop. Primers THR (forward) and 651 (reverse) were used for initial amplification of entire ~1300 bp. Internal primers 007 (forward), DL3 (reverse), and DL1 (reverse) were used for Sanger sequencing (Shaffer & McKnight, 1996).

### *Phylogenetic Reconstructions* for Mitochondrial Haplotypes

Phylogenetic reconstructions were estimated for all unique mitochondrial D-loop haplotypes and an additional 53 *A. barbouri*, 83 unisexual Ambystomatids and 27 *A. texanum* sequences obtained from Genbank. Both maximum likelihood (ML) and Bayesian optimality criteria were used for phylogenetic analyses. Maximum-likelihood analyses were performed using the software RAxML (Stamatakis 2014) on the CIPRES Science Gateway (Miller et al. 2010) under the GTR+G model. Nodal support was estimated using rapid bootstrapping (1000 replicates). Bayesian phylogenetic reconstructions were performed using MrBayes 3.2.1 (Huelsenbeck & Ronquist 2001) also on the CIPRES Science Gateway. The best model of substitution was selected by Modeltest (Posada & Crandall 1998), implemented in MEGA X (Kumar et al. 2016) using Bayesian information criterion (BIC). The Markov chain Monte Carlo (MCMC) algorithm ran for 10,000,000 generations, sampling every 1,000 generations. Two independent runs were performed and the resulting trees were combined after the deletion of a burnin (25%). A majority-rule consensus tree was generated and nodal support was estimated by posterior probabilities.

### GBS Library Preparation and Sequencing

A total of 227 individuals were included for GBS sequencing including 201 individuals from 12 populations of *A. barbouri* in Tennessee, 17 individuals from five populations of *A. barbouri* in Kentucky, five individuals from two populations of *A. texanum*, and two sampled outgroups *A. talpoideum* (N=1) and *A. mabeei* (N=1; Table 1). Genotyping-by-Sequencing libraries were prepared using the restriction enzyme ApeKI following the protocol of Elshire et al. (2011). Genomic DNA was quantified using Quant-iT Picogreen dsDNA Assay Kit (Thermo Fisher Scientific), and all samples were standardized to between 5-6.5 ng/uL (50-65 ng total genomic DNA). Extracted DNA was digested with the restriction enzyme ApeKI. Adaptors containing PCR binding sites and individual barcodes were ligated onto digested DNA. Barcoded DNA was pooled, and PCR was amplified using primers that bind to the ligated adaptors (see Elshire et al. 2011 for primer sequences). The resulting PCR products were cleaned using the Qiagen PCR purification kit and then cleaned again using the AxyPrep Mag PCR Clean-up kit (Axygen, Big Flats, New York, USA). The distribution of the fragment size in the PCR product was determined using an Agilent 2100 BioAnalyzer (Agilent Technologies Inc., Santa Clara, CA, USA). Barcoded libraries were sequenced using the Illumina NextSeq (Illumina Inc., San Diego, CA, USA) with a 75 bp single end read chemistry.

### SNP Discovery and Filtering

The *Stacks* program *process_radtags* was used to filter and demultiplex raw reads based on barcoded sequences (Catchen 2011). We used the seven-step de novo clustering pipeline ipyrad v. 3.5 (Eaton 2014) to generate and filter SNP datasets used in downstream analyses. Quality filtering of raw sequence reads converted bases with Phred scores <33 to Ns, reads with more than 5 Ns were removed. Reads were clustered using a sequence similarity threshold of 90% both within and between sampled individuals, with a minimum read depth of six. Individuals with fewer than 500,000 reads were excluded from downstream analyses. Loci with observed heterozygosity (H_o_) greater than 0.5 were removed to filter out possible paralogs. The final SNP dataset was then filtered to remove loci deviating from Hardy-Weinberg equilibrium (p < 0.05), loci genotyped in less than 60% of individuals, and SNPs with a minor allele frequency (MAF) less than 0.01. Only one SNP per tag was retained per locus.

### Population Genetic Variation and Effective Population Size

Estimates of within population genetic variation for 12 Tennessee *A. barbouri* populations were estimated using the R-package DiveRsity v. 1.9.9 (Keenan et al. 2013). Summary statistics were estimated separately for each population and included the proportion of SNPs that were polymorphic within each population (*P*), allelic richness (*A_R_*), and observed and expected heterozygosity (*H_o_* and *H_e_*). Estimates of *P*, *A_R_*, *H_o_*, and *H_e_* were estimated for the entire SNP dataset and for the subset of SNPs that were polymorphic within each population.

Effective population size (N_E_) was estimated for Tennessee *A. barbouri* populations using both the linkage disequilibrium (LD) method and the heterozygote-excess method (Zhdanova & Pudovkin 2008) as implemented in the program N_E_Estimator v2 (Do et al. 2014). Populations with sample sizes of less than 20 individuals were excluded from analyses as these datasets did not provide enough signal for reliable N_E_ estimation (Nunziata & Weisrock 2018). The minor allele frequency (MAF) parameter was set at 0.01 and 95% confidence intervals were estimated using the parametric chi-squared method.

### Population Structure

Population pairwise F_ST_ values were estimated for the all *A. barbouri* populations (Weir & Cockerman 1984). Hierarchical partitioning of genetic variation across Tennessee populations of *A. barbouri* was examined using Analysis of Molecular Variances (AMOVA) as implemented by Arlequin 3.5 (Excoffier & Lischer 2010, Excoffier et al. 1992). Results from phylogenetic reconstructions based on mitochondrial D-loop haplotypes were used to generate hypotheses regarding higher-level structuring of populations. Significance of variance components was determined using 1000 permutations.

The optimal number of genetic clusters (K) based on genomic SNPs was estimated using both a multivariate approach and a Bayesian-based assignment method. Discriminant Analysis of Principal Component (DAPC) was performed using the ‘adegenet’ package in *R* (Jombart & Ahmed 2011, Jombart & Collins 2015). The *find.clusters* function was first used to identify the optimal value of *K* based on a Bayesian Information Criterion (BIC) process. The optimal number of principal components was determined using the function *a.score.* The optimal K was then used to perform a DAPC analysis to describe the relationship between the genetic clusters and individual membership probabilities were assessed using the function *compoplot*. DAPC analysis was performed on three hierarchical datasets; these included 1) 12 populations of *A. barbouri* in Tennessee, 2) all sampled Tennessee and Kentucky *A. barbouri* populations and 3) all *A. barbouri* populations and four *A. texanum* individuals from AAFB.

Bayesian assignment tests were performed on Tennessee *A. barbouri* populations using the program STRUCTURE 2.3.4 (Pritchard et al. 2003). Kentucky *A. barbouri* populations were not included in assignment analyses due to small sample sizes at these sites. Values of *K* (number of populations) ranged from 1 to 12 populations with 20 replicate runs per value of *K*; MCMC simulations were performed for a burn-in of 250,000 iterations and an additional 1,000,000 iterations were retained for the final analysis. Results were summarized using the software package CLUMPAK (Kopelman et al. 2015). The optimal number of groups (*K*) was determined using the Delta-K method (Evanno et al. 2005) as implemented in STRUCTURE HARVESTER v0.6.94 application (Earl & VonHoldt 2012). The Distruct (1.1) application in CLUMPAK (Kopelman et al. 2015) was used to produce the final barplots.

### Multispecies Coalescent using BPP

A multispecies coalescent (MSC) model was used to examine taxonomic boundaries, estimate species trees, and estimate population divergence times (*t* = years before present) as implemented in the software BPP v. 4.4 (Yang 2015). Input datasets were formatted as full-length sequence alignments from our GBS sequencing and included all *A. barbouri* and *A. texanum* populations. Loci represented by fewer than 100 individuals were removed from the analysis. Joint species-delimitation and species-tree reconstructions were performed using the A11 model, where individuals were assigned to sampled populations. Run parameters were as follows: uniform rooted trees were used as the species model prior, the theta prior (θ = 4N_Eμ_) was assigned a gamma distribution with α=3 and β=0.04 and the prior for tau (τ=tμ) was gamma distributed with α=3 and β=0.2. Three independent MCMC runs were implemented, each with a burn-in of 5,000, a sample frequency of 10 and a total of 50,000 iterations. Species delimitation and species-tree topologies with the highest posterior probabilities from the A11 runs were used as input for the A00 analyses. The A00 model estimates divergence time parameters and long-term NE, assuming fixed species assignments and a fixed species tree. The priors for A00 were the same as priors used for the A11 model. Three independent runs were also performed for the A00 analysis, each with a burn-in of 5,000, a sample frequency of 10 and a total of 50,000 iterations. MCMC chains were pooled and absolute estimates of τ were converted to years before present (YBP) in the BPPR using the genome wide mutation rate estimate for the vertebrate nuclear genome where μ= 1.21 х10^−9^ (Allio et al. 2019, Reis & Yang 2019).

## Results

### Mitochondrial Sequencing

Sanger sequencing of the mitochondrial D-loop resulted in an ~1100 bp sequence alignment in the 81 individuals selected for sequencing. A total of 60 unique haplotypes were identified from the newly sequenced individuals, 39 haplotypes were identified from *A. barbouri* populations sampled in Tennessee (See Supplement 1 for Genbank Accessions) and 15 haplotypes were identified from populations sampled in Kentucky. An additional six haplotypes were sequenced from outgroups. The final alignment contained 126 variable sites (excluding outgroups); of these, 120 sites were parsimony informative. No haplotypes were shared across multiple populations.

### Phylogenetic Reconstructions

Both Bayesian and ML phylogenetic reconstructions of mitochondrial D-loop haplotypes identified six major clades across *A. barbouri* and *A. texanum* haplotypes (Fig. 3). *Clade I* included all haplotypes belonging to the unisexual lineage of *A. barbouri* with 100% posterior probability (pp) and 100% bootstrap support (bs) for Bayesian and ML analysis, respectively. In *Clade II*, *A. barbouri* from Wilson Co. (populations W1, W3, and W4) and Davidson Co. (population D3) formed a monophyletic clade that also includes Kentucky *A. barbouri* populations from southwest of the Kentucky River (pp = 100, bs = 95%). *Clade I* and *Clade II* were recovered as sister clades in both analyses with strong support (pp = 100, bs = 99%). *Clade III* included all *A. texanum* individuals (pp = 99, bs = 71) with the exception of a single *A. texanum* haplotype obtained from Genbank (ID EU980569, collected from Lawrence KA); this sequence is recovered as basal to *Clade III* in both ML and Bayesian analyses. *Clade IV* groups with *Clade III* and included Tennessee *A. barbouri* individuals from Rutherford Co. (populations R1, R7, and R9), Bedford Co. (B6) and Williamson Co.; pp = 100, bs = 71%). *Clade V* included all Kentucky individuals from populations N_E_ of the Kentucky River, Ohio and Indiana (pp = 100, bs = 86%). Finally, *Clade VI* is sister to *Clade V* and included all Tennessee *A. barbouri* from Sumner Co. (populations S2, S5, S7, and S8), a single haplotype from Wilson population W3 and three haplotypes sequenced from Rutherford population R7 (pp = 100, bs = 99%). On a larger scale, all *A. barbouri* and *A. texanum* are part of the same well-supported monophyletic group in both analyses (pp = 100, bs = 100%), such that all *A. texanum* haplotypes are nested within *A. barbouri*.

**Figure 3.**
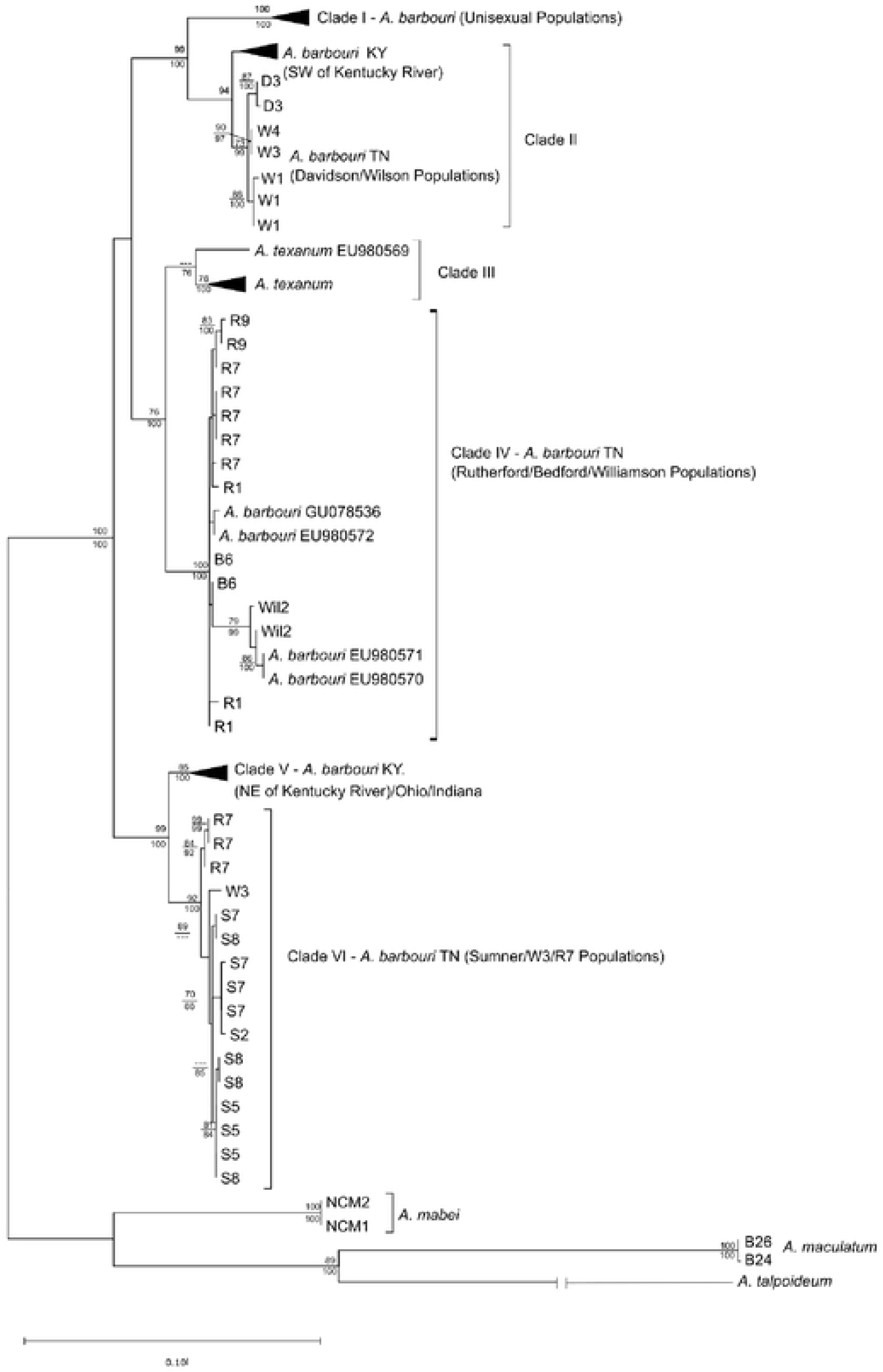
Maximum likelihood phylogenetic reconstrnctions of unique mitochondrial D-loop haplotypes from *Ambysloma barbouri* and *A. texanum* under a GTRCAT model of evolution as perfonned by RaxML Bootstrnp support values above branches are shown for nodes with 70% support or greater Values below nodes indicate posterior probabilities from Bayesian reconstructions under a T92+G model of sequence evolution as perfonned by MrBayes. Asterisks denote accession numbers for sequences downloaded from Genbank.

### GBS sequencing

The average number of retained sequence reads per individual generated from sequencing of GBS libraries was 5,780,366; 220 individuals were retained after removal of low coverage individuals (defined as < 500,000 total reads). A total of 1,558,844 loci were recovered from the de novo assembly in ipyrad and 440,629 loci were retained after filtering.

Additional filtering for SNPs (i.e. filtering for low representation, HWE, MAF, and max heterozygosity) recovered 1,169 SNPs for the dataset that included only Tennessee *A. barbouri*, 516 SNPs in the dataset that included all sampled *A. barbouri* populations in Tennessee and Kentucky, and 500 SNPs in the dataset that included all *A. barbouri* and *A. texanum* populations.

### Within Population Genetic Variation

Much of the genetic variation within our SNP dataset was in the form of fixed differences between populations (Table 3). The proportion of SNPs that were polymorphic within populations ranged from 0.045 (W3/Wil2) to 0.267 (S7). When only polymorphic loci were considered, the Will2 population had the highest estimates of Ho and He; however, this population had a low proportion of polymorphic SNPs overall. Rutherford County populations (R1/R7) had the lowest estimates of H_o_ and H_e_, when considering only polymorphic loci and for the entire SNP dataset.

**Table 3.**
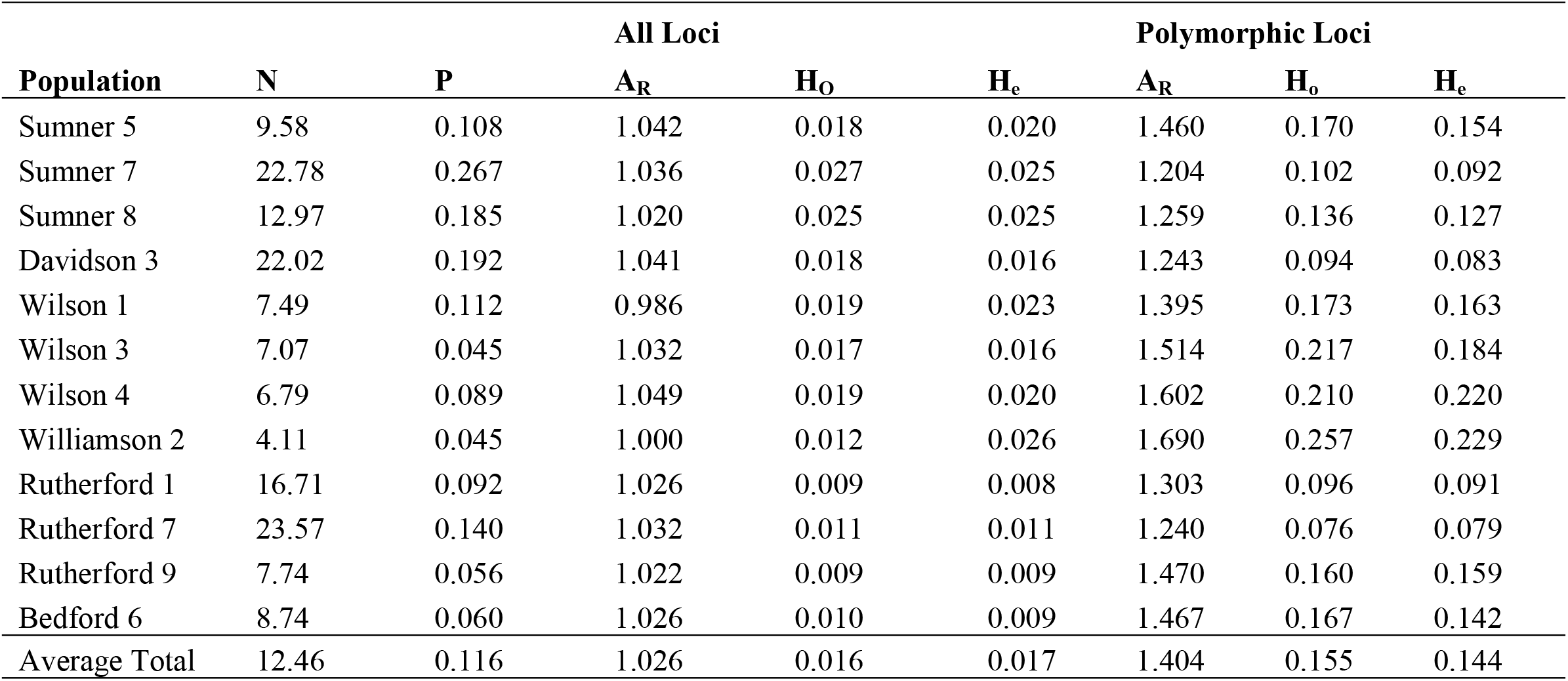
Standard measures of genetic diversity for 12 populations of *Ambystoma barbouri* in Tennessee based 586 SNP loci. Summary statistics include the average number of individuals genotyped per locus (*N*) and the proportion of SNPs that were polymorphic within each population (*P*). Allelic richness (A_R_), observed heterozygosity (H_O_), and expected heterozygosity (H_e_) were calculated for all loci and again for only those loci that were polymorphic within each population.

### Effective Population Size

Estimates of effective population size (*N_E_*) were generated for five populations with adequate sampling (*N* ≥ 20), including S7, S8, R1, R7, and D3 (Table 4). Results from LD analyses ranged from *N_E_* =15 (R1) to *N_E_* = 108 (108) (S8); the upper confidence interval for S8 was ∞, indicating low signal for this dataset. Results from the heterozygote excess method were inconclusive (∞) for two of the five analyzed populations. Estimates from the remaining three populations ranged from 24 (D3) to 406 (S7); however, CIs included ∞ for two of these estimates (S7 and S8).

**Table 4.** Estimates of effective population sizes (N_E_) for five populations of *Ambystoma barbouri* in Tennessee based on the Linkage Disequilibrium (LD) method and the Heterozygosity Excess Method as performed by N_e_Estimator. The 95% confidence intervals were estimated by the parametric chi-squared method.

| Population | $N$ | Linkage Disequilibrium | | | Heterozygote Excess | | |
| --- | --- | --- | --- | --- | --- | --- | --- |
| | | $N_E$ | CI Lower | CI Upper | $N_E$ | CI Lower | CI Upper |
| Sumner 7 | 35 | 84.5 | 51.5 | 207.2 | 406.4 | 20.7 | ∞ |
| Sumner 8 | 21 | 107.7 | 28.8 | ∞ | 192.7 | 18.8 | ∞ |
| Davidson 3 | 29 | 52.1 | 33.2 | 106.4 | 24.0 | 13.0 | 182.8 |
| Rutherford 1 | 20 | 14.8 | 8.7 | 30.1 | ∞ | 14.6 | ∞ |
| Rutherford 7 | 28 | 30.7 | 20.9 | 51.6 | ∞ | 22.8 | ∞ |

### Population Structure

Pairwise F_ST_ estimates for *A. barbouri* populations within Tennessee were geographically structured in a hierarchical fashion (Table 5). Overall, F_ST_ values averaged 0.375 and ranged from no differentiation (FST= 0) to highly differentiated (F_ST_ = 0.725, S5/R1). In general, populations north of the Cumberland River (S2, S5, S7, and S8) showed evidence of long-term isolation from populations south of the Cumberland. Pairwise F_ST_ estimates comparing populations north and south of the Cumberland averaged 0.637, while F_ST_ estimates among Tennessee populations north of the Cumberland averaged only 0.043. Populations south of the Cumberland were further structured; populations in Wilson and Davidson Counties (W1, W3, W4, and D3) were diverged from populations further south in Williamson, Rutherford and Bedford counties. Estimates of pairwise F_ST_ comparing populations within Wilson/Davidson averaged 0.121 and within Williamson, Rutherford and Bedford Counties averaged 0.070, while between group pairwise F_ST_ values were much higher, averaging 0.300.

**Table 5.** Population pairwise F_ST_ estimates averaged across 584 SNP loci for 12 populations of *Ambystoma barbouri* in Tennessee (below diagonal) and significance of F_ST_ estimates (p-values, below diagonal) estimated from nonparametric permutations of SNP genotypes (100 permutations) as performed by the software Arlequin. ** indicates *p* < 0.001.

| Populations | 1 | 2 | 3 | 4 | 5 | 6 | 7 | 8 | 9 | 10 | 11 | 12 |
| --- | --- | --- | --- | --- | --- | --- | --- | --- | --- | --- | --- | --- |
| 1. Sumner 5 | --- |  |  |  |  |  |  |  |  |  |  |  |
| 2. Sumner 7 | 0.053 |  |  |  |  |  |  |  |  |  |  |  |
| 3. Sumner 8 | 0.022 | 0.055 |  |  |  |  |  |  |  |  |  |  |
| 4. Davidson 3 | 0.650 | 0.614 | 0.647 |  |  |  |  |  |  |  |  |  |
| 5. Wilson 1 | 0.670 | 0.616 | 0.641 | 0.207 |  |  |  |  |  |  |  |  |
| 6. Wilson 3 | 0.660 | 0.590 | 0.622 | 0.055 | 0.104 |  |  |  |  |  |  |  |
| 7. Wilson 4 | 0.581 | 0.527 | 0.558 | 0.111 | 0.157 | 0.092 |  |  |  |  |  |  |
| 8. Williamson 2 | 0.669 | 0.568 | 0.602 | 0.203 | 0.238 | 0.278 | 0.167 |  |  |  |  |  |
| 9. Rutherford 1 | 0.725 | 0.638 | 0.698 | 0.288 | 0.408 | 0.399 | 0.320 | 0.031 |  |  |  |  |
| 10. Rutherford 7 | 0.704 | 0.637 | 0.687 | 0.276 | 0.234 | 0.322 | 0.239 | 0.077 | 0.127 |  |  |  |
| 11. Rutherford 9 | 0.695 | 0.601 | 0.651 | 0.285 | 0.356 | 0.370 | 0.276 | 0.039 | 0.021 | 0.113 |  |  |
| 12. Bedford 6 | 0.698 | 0.600 | 0.655 | 0.280 | 0.386 | 0.385 | 0.292 | 0.000 | 0.051 | 0.157 | 0.085 | --- |

The three monophyletic clades of Tennessee *A. barbouri* recovered from mitochondrial phylogenetic reconstructions were further examined using genome-wide SNP genotypes in an AMOVA framework (Table 6). The majority of variance in SNP genotypes could be attributed to differences between the three clades (63.64%). Differences between populations within clades only accounted for 2.09% of the total molecular variance. Differences between individuals accounted for the remaining 34.27% of variance in SNP genotypes.

**Table 6.** Results of hierarchical analyses of molecular variation (AMOVA) for the SNP dataset from 12 populations of *A. barbouri* in Tennessee. Assignment to mitochondrial clades are as follows: Clade II (S5, S7, and S8), Clade IV (D3, W1, W3, and W4), and Clade III (Wil2, R1, R7, R9, B6). Asterisks indicate significance of Φ statistics based on 1000 permutations in Arlequin.

| Source of variation | DF | Sum of squares | Variance components | Percentage of variation | Φ-Statistics |
| --- | --- | --- | --- | --- | --- |
| Among drainages | 2 | 614.5 | 2.30 | 63.60 | $\phi_{CT} = 0.636^*$ |
| Among populations within drainages | 9 | 31.24 | 0.08 | 2.09 | $\phi_{SC} = 0.058^*$ |
| Within populations | 384 | 476.24 | 1.24 | 44.23 | $\phi_{ST} = 0.657^*$ |
| Total | 395 | 1,121.97 | 3.62 |  |  |

Results of BIC for DAPC analysis suggested that three genetic clusters (*K*=3) best represented genetic variation among populations sampled within Tennessee (Fig. 4A, Supp 2). Membership of each cluster was geographically partitioned and consisted of a northern cluster that included populations from Sumner County, a central cluster comprised of individuals from Wilson and Davidson Counties, and a southern cluster that represented Bedford, Rutherford, and Williamson County populations. Membership probabilities and DAPC plots indicated some degree of admixture between the central and southern clusters, but no admixture within the northern cluster (Supplement 2). A separate analysis that included Tennessee *A. barbouri* and the five sampled populations from Kentucky also resulted in three genetic clusters (Fig 4B).

**Figure 4.**
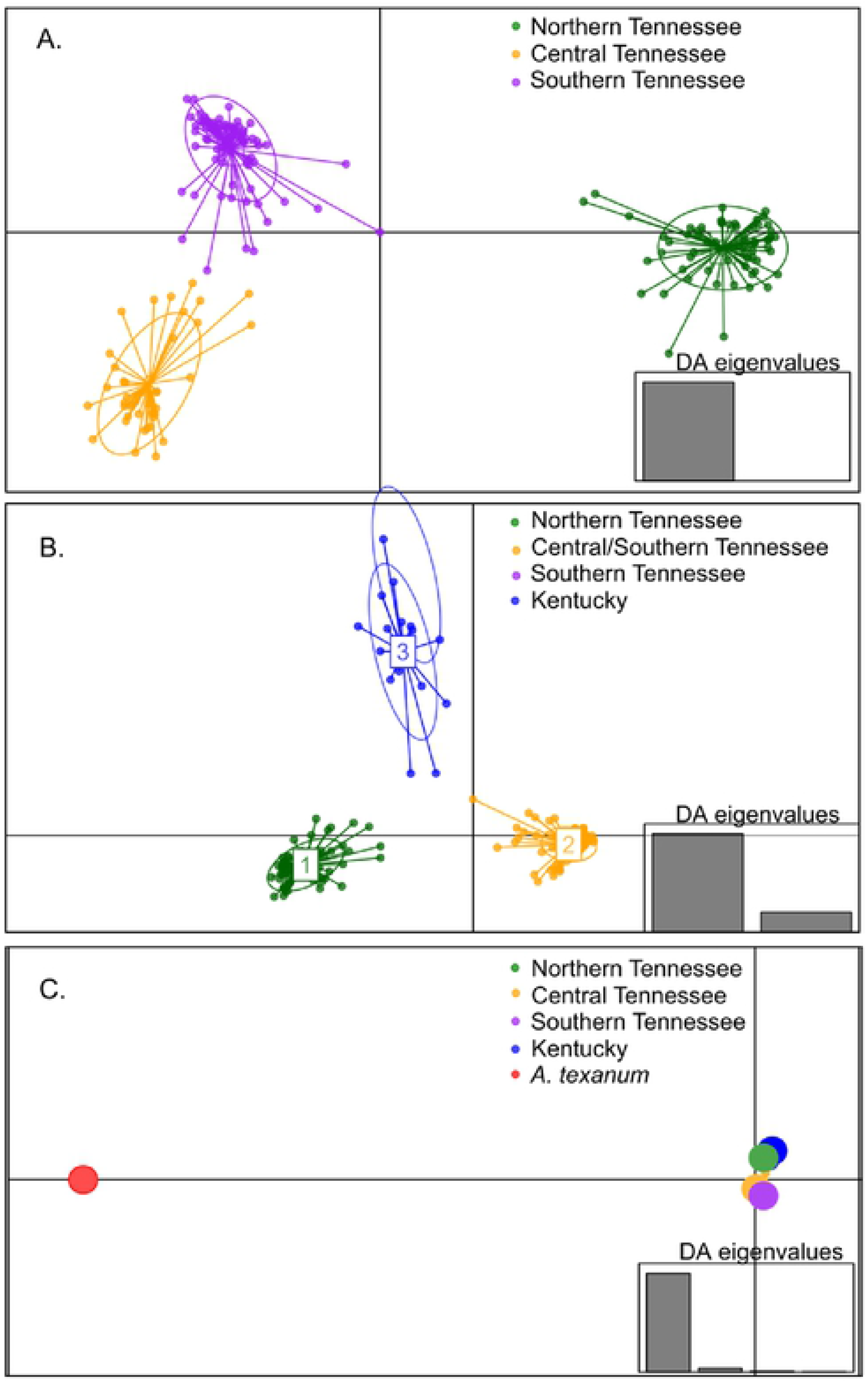
Discriminant Analysis of Principal Components (DAPC) results based on SNP genotypes A.) DAPC plot for *A. barbouri* (*N* = 198) from Tennessee Analysis assumed a value of K = 3 as the optimized by BIC analysis B) DAPC plot for *A. barbouri* from Tennessee (N = 198) and also including samples from five populations of *A barbouri* in Kentucky *(N* = 17). Analysis was performed using K =3 as the optimal number of genetic clusters based on BIC analysis. C) DAPC plot for *A. barbouri* from Tennessee. Kentucky. and also two populations of *A. texanum* (N = 4). Analysis was pcrfonned using K = 5 as the optimal number of genetic clusters according to the BICanalysis.

Kentucky populations of *A. barbouri* were isolated from populations in Tennessee. Central and southern populations were assigned to the same cluster and populations in Sumner county were assigned to the third cluster. When *A. texanum* individuals were included in a third analysis, results of the BIC analysis indicated the number of genetic clusters was five (Fig. 4C). Tennessee populations were assigned to northern, central, and southern clusters and scatter plots showed Tennessee populations to be tightly grouped with the fourth cluster that included all Kentucky populations. All *A. texanum* individuals formed the fifth cluster that was well separated from all *A. barbouri* individuals.

Analysis of Bayesian assignment tests for Tennessee populations of *A. barbouri* were consistent with results from DAPC analysis that suggested the presence of three geographically partitioned genetic clusters within the state (Fig. 5). The optimal value of *K*, as determined by the Delta K method, indicated four genetic clusters; however, increasing K from 2 to 3 did not change population assignments. Assignment plots for K=2-3 separated northern populations in Sumner County from central and southern populations. When K was increased to four, populations from Davidson and Wilson Counties were separated from southern populations in Bedford, Rutherford and Williamson Counties. Increasing the value of K to 5 increased the level of admixture, but did not result in geographically meaningful partitions.

**Figure 5.**
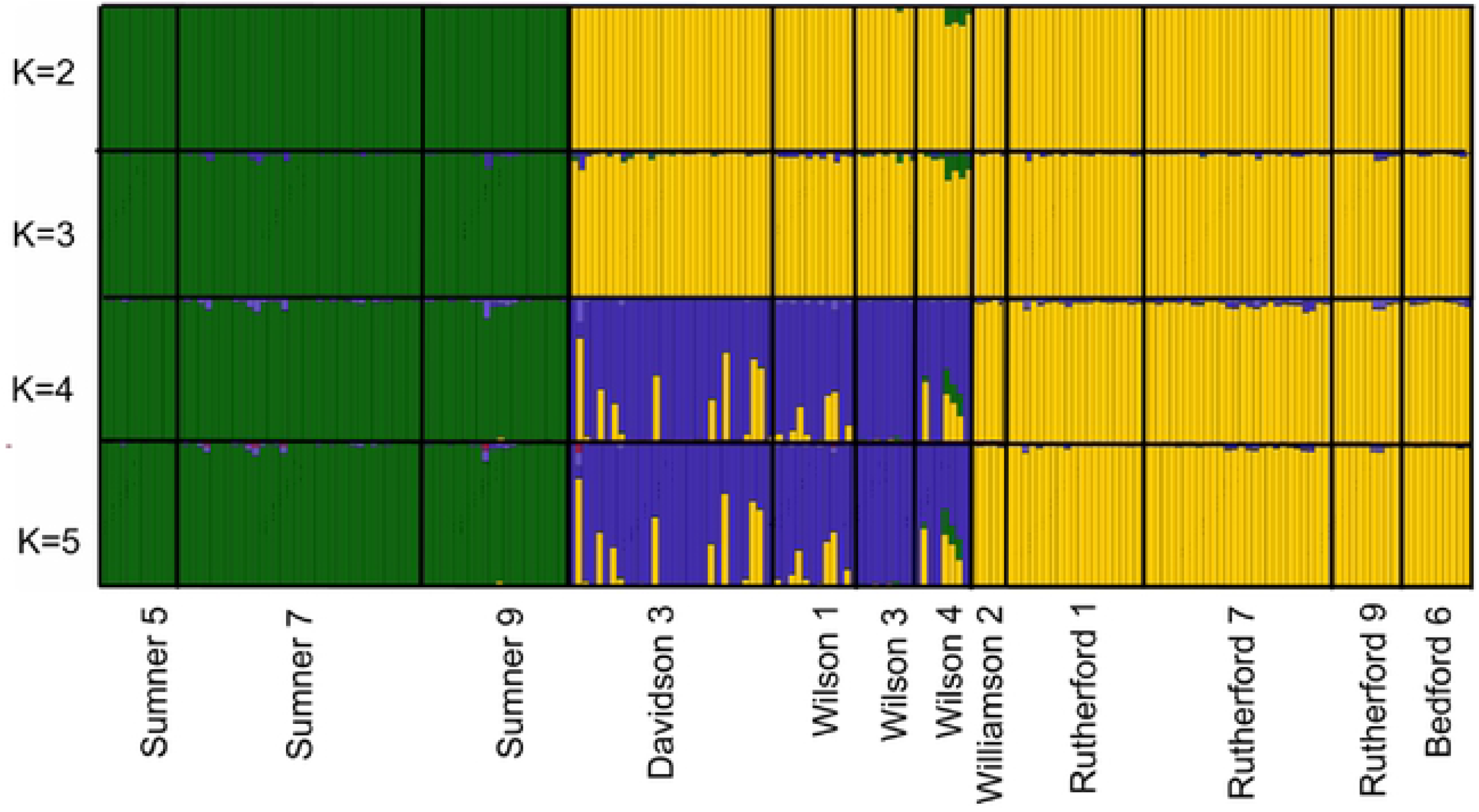
Results of Bayesian Assigmnent tests based on SNPs generated from GBS sequencing. Baiplots indicate individual assignment probabilities for samples from 12 *A. barbouri* populations in Tem1essee (*K* = 2-5).

### Multispecies Coalescent Analyses

A total of 988 loci were retained for the MSC analyses performed by BPP. All three A11 runs recovered the same taxonomic groups and species tree topology with the highest posterior probability. Out of 18 sampled populations, 14 populations were genetically distinct. Within Tennessee, *A. texanum* and all *A. barbouri* populations, with the exception of R1/R7, were identified as distinct groups (Fig. 6). Kentucky *A. barbouri* populations were split into two groups; populations FC, JC, RR, and SL were grouped together and the SW populations was its own genetic group. The separation of individual populations in this analysis does not indicate that each population warrants recognition at the species-level as simulations have shown that BPP will split groups at the population-level when many loci are analyzed (Leach et al. 2019).

**Figure 6.**
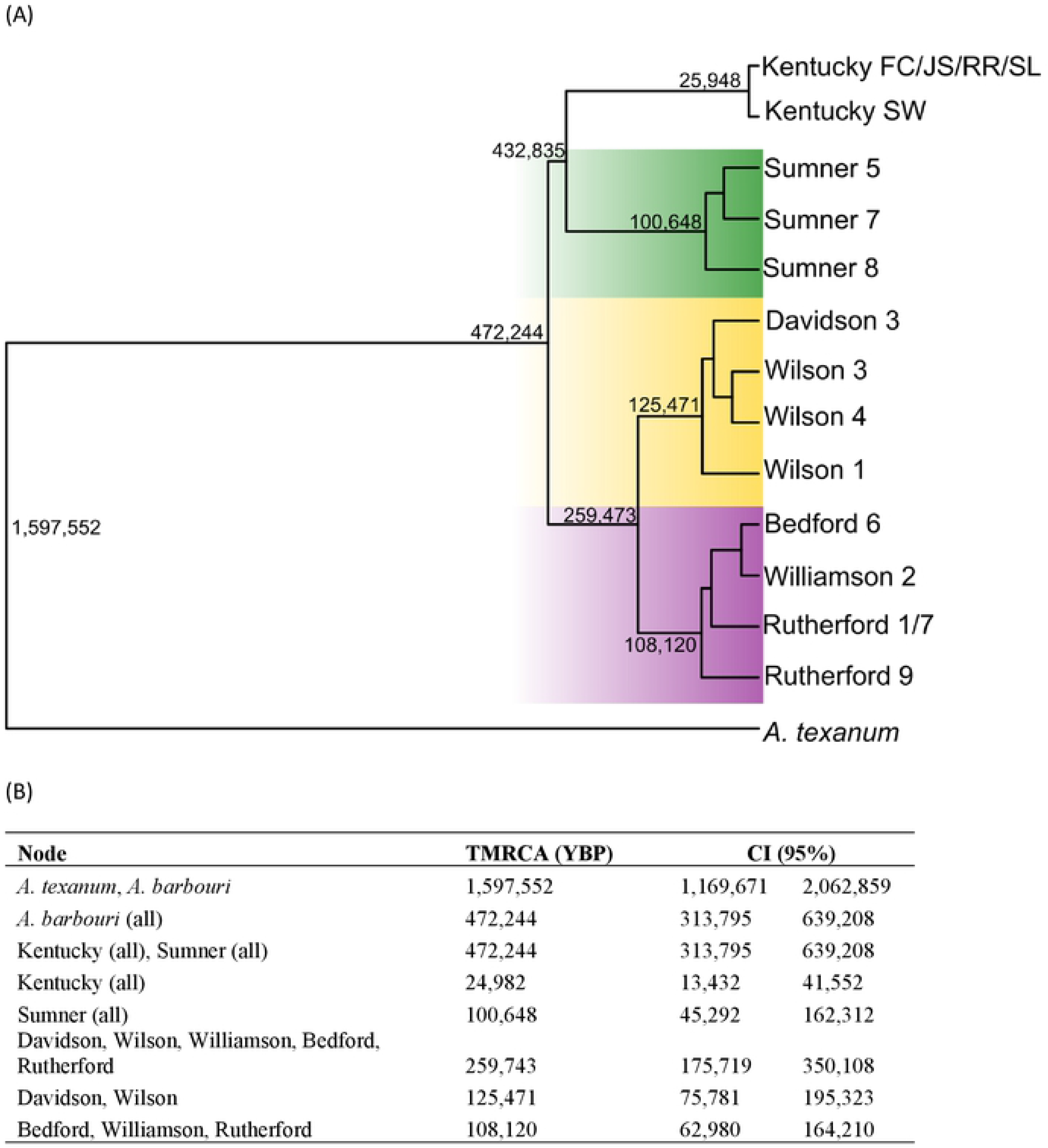
Species-tree (A) and divergence time estimates (B) from Bayesian multispecies coalescent method as implemented in the software BPP Numbers at nodes also indicate divergence lime estimates *(t)*. Divergence time parameter twas converted 10 time in years before present (YBP) in the BPPR statistical package in R studio. Shading in tree indicates ESU assignments as follows: Northern ESU (green), Central ESU (Yellow) and Southern ESU (Purple).

The A11 species tree reconstruction with the highest posterior identified the same Tennessee clades recovered in our mitochondrial gene trees with the exception of the placement of *A. texanum*. In the species tree, *A. texanum* was basal to all populations of *A. barbouri* in Tennessee and Kentucky. The estimated divergence time (A00 analysis) between *A. texanum* and *A. barbouri* was 1.6 million YBP. All Sumner *A. barbouri* formed a monophyletic group with Kentucky populations as a monophyletic sister group. Divergence time estimates indicated that Sumner populations and Kentucky populations shared a common ancestor < 500,000 YBP. All *A. barbouri* populations south of the Cumberland River were monophyletic (TMRCA 260,000 YBP) and were further split into two groups that corresponded to clades recovered in the mitochondrial gene tree. Davidson and Wilson populations were recovered together in the first group (mitochondrial Clade II) and Rutherford, Bedford, and Williamson populations (mitochondrial Clade IV) formed the second group.

## Discussion

Patterns of genomic variation and taxonomic relationships identified here have important implications for developing management strategies aimed at the long-term conservation of *A. barbouri*. Despite the documented decline of *A. barbouri* populations in the Nashville Basin, results from GBS-derived SNP genotyping indicate that estimates of genetic variation in extant populations of Tennessee *A. barbouri* are similar to SNP-based datasets examined from other ambystomatids. Partitioning of genetic variation between Tennessee populations suggests that hydrogeography of the Nashville Basin has shaped patterns of gene flow. Both mtDNA and genomic SNP genotypes showed similar patterns with respect to population structure within the state of Tennessee that should be used to inform the designation of units of conservation. Our results also reveal a complex history of Tennessee *A. barbouri* with more northern populations and with its sister species *A. texanum*. Despite marked differences in phylogenetic reconstructions based on mtDNA sequencing and nuclear genotypes, collectively these results indicate that Tennessee is genetically unique from Northern *A. barbouri* and *A. texanum.*

### Genetic Variation within Populations

Loss of genetic variation resulting from rapid population decline threatens the long-term success of conservation efforts due to loss of adaptive potential and fixation of deleterious alleles (Kardos et al. 2021). For *A. barbouri* populations in the Nashville Basin, maintenance of adaptive genetic variation is particularly critical as these populations are faced with habitat alteration by urbanization in addition to climate change (Schmidt et al. 2021). Evaluating the genetic health of populations is valuable for management and conservation planning; however, the interpretation of genetic variation estimates from SNP data is challenging due to the limited number of comparable studies utilizing reduced-representation methods in salamanders. Heterozygosity estimates obtained here are similar to estimates obtained in the handful of published SNP-based genetic studies. In a recent survey of SNPs from ddRAD sequencing, estimates of H_O_ ranged from 0.165 to 0.269 for populations of the mole salamander (*A. talpoideum*) and were slightly higher for populations of the marbled salamander (*A. opacum*; 0.207 - 0.298; Nunziata et al. 2017). Another SNP-based survey examined the effects of land use on genetic variation in the northern two-lined salamander (*Eurycea bislineata*). Fusco et al. (2020) reported nearly identical estimates of H_O_ from urban, suburban and rural salamander populations (H_O_= 0.265, 0.278 and 0.275, respectively) and concluded that genetic variation had been maintained, despite habitat disturbance. However, estimates of heterozygosity may not be directly comparable across independent studies as filtering parameters in bioinformatic pipelines can influence population genetic summary statistics. Stringent filtering criteria may preferentially retain loci in conserved regions of the genome and downwardly bias estimates of genetic variation (Huang & Knowles 2014). There have been previous population genetic studies in *A. texanum* and *A. barbouri* populations based on microsatellite markers. Micheletti and Storfer (2017) estimated genetic variation at 11 microsatellite loci in 76 populations of *A. barbouri* distributed throughout Kentucky, Ohio, and Indiana; the average H_e_ from their study ranged from 0.67 – 0.81, respectively. Few conclusions can be made by comparing results from these prior microsatellites studies to our results based on SNP genotypes; microsatellites are known to have a much higher rate of mutation than nucleotide substitutions, increasing the expected amount of allelic diversity for a given population size (Haasl & Payseur 2011).

### Effective population size

A reduction in estimates of contemporary effective population size (N_E_) may also signal a decline in the genetic health of at-risk populations. Our estimates of N_E_ were low (NE from the LD method averaged 58.0 across five populations), but were not unusual for LD-derived estimates of N_E_ in salamanders. Published estimates of N_E_ reported for amphibians have typically been under 100 (Jehle & Arntzen 2002). Effective population size estimates reported here were similar to values reported for the congeneric endangered California tiger salamander (*A. californiense*; N_E_ values of 11-64; Wang et al. 2011), and for the long-toed salamander (*A. macrodactylum*; N_E_ = 23-207; Funk et al. 1999). In *A. macrodactylum*, Savage et al. (2010) estimated N_E_ for 47 breeding populations and more than half of these estimates were less than 50. Life history factors may provide some explanation for low N_E_ in this group as salamanders often exhibit high variance in reproductive success and larval survival within populations. Variance in reproductive success will increase relatedness among individuals in a population, inflating linkage disequilibrium across loci and reducing N_E_.

Obtaining robust estimates of N_E_ and other demographic parameters in at-risk species can be challenging due to the sample sizes required for LD detection. Many threatened taxa are inherently rare and prohibitively difficult to sample in large numbers. For robust estimation of N_E_, it is recommended that sample sizes be greater than 30 for most systems (. When the number of sampled individuals is much smaller than the effective size, LD-based N_E_ estimates can be downwardly biased and confidence intervals can be large (or infinite) due to inadequate signal in the dataset (Waples & Do 2010). We limited our N_E_ analysis to populations with sample sizes > 20 individuals, which left only five populations for N_E_ estimation. The infinite upper confidence interval for our N_E_ estimate in population Sumner 8 was likely due to an inadequate sample size for this population (*N*=21).

### Phylogenetic Reconstructions and Population Structure

Mitochondrial gene-tree reconstructions suggest a complex biogeographic history for *A. barbouri* in Tennessee that involves populations of unisexual Ambystomatids, *A. texanum* and populations of *A. barbouri* from the northern core region (Kentucky, Indiana, and Ohio). Our results support findings by Bogart et al. (2007) demonstrating a monophyletic relationship of unisexual Ambystomatid mtDNA haplotypes nested within *A. barbouri*. The phylogenetic reconstruction shown here further details this relationship whereby unisexual Ambystomatids likely shared a maternal common ancestor with *A. barbouri* populations in Kentucky and populations from Middle Tennessee. This pattern, coupled with phylogenies based on nuclear genes, points to a history of repeated hybridization events initiated by an *A. barbouri* maternal ancestor in the southern portion of its range.

With regards to their mitochondrial lineage, *A. texanum* is nested within present-day *A. barbouri,* rendering *A. barbouri* paraphyletic. This phylogenetic pattern contradicts the scenario proposed by Kraus and Petranka (1989), that stream-dwelling *A. barbouri* descended from the more widespread pond-breeding *A. texanum*. Species-tree reconstructions based on genome-wide SNP data are incongruent with relationships recovered from the mitochondrial trees. This species-tree topology adheres to conventional taxonomic and geographical boundaries (Fig. 6), where *A. texanum* is a sister-species to all *A. barbouri* populations. The TMRCA for *A. barbouri* and *A. texanum* based on SNP genotypes was estimated at ~1.6 million YBP making this a relatively recent split between these ecologically distinct species (Vences et al. 2007).

Incongruence between mitochondrial gene trees and reconstruction based on nuclear data are not uncommon and are often attributed to ancestral lineage sorting, introgression, and/or sex-biased dispersal (Nichols 2001, Toews & Brelsford 2012). Determining the cause of mitonuclear discordance would require additional sampling across the distributions of both *A. barbouri* and *A. texanum;* however, it is relevant to note that recent studies have demonstrated that hybridization is common across salamander lineages (including the genus *Ambystoma*) and may facilitate rapid diversification (Patton et al. 2020).

The hydrogeography of the Central Basin, together with cyclic glacial movements during the Pleistocene, may have shaped contemporary patterns of genetic variation in Tennessee populations of *A. barbouri*. Mitochondrial-based phylogenetic reconstructions and partitioning of genetic variation at SNP loci (including assignment tests, DAPC analyses, AMOVA, and MSC reconstructions) identify the same three genetically distinct groups in Tennessee; these include a northern cluster, a central cluster, and a southern cluster. The Cumberland River appears to have served as a major barrier to gene flow between the northern and the central/southern clusters. The central and southern clusters both occur south of the Cumberland River and are divided by smaller regional drainage patterns. Individuals in the northern and central clusters occupy the Old Hickory Lake and Lower Stones River watersheds, respectively. The central cluster is bordered by the Cumberland River to the North and by the East Fork of the Stones River to the east. The southern cluster includes populations from three watersheds including the West Fork Stones River Watershed, Upper Duck River Watershed, and Mill Creek Watershed; the southern cluster (with the exception of the Wil2 population) is bordered by the Stones River to the north and by the Duck River to the south.

Mitochondrial-based phylogenetic reconstructions revealed that Tennessee *A. barbouri* populations are not monophyletic with respect to northern populations, which may reflect a history of repeated range expansions from Tennessee into the northern end of *A. barbouri’*s present day distribution. The estimated timing of divergence between the three Tennessee clusters falls within the mid-late Pleistocene. Glacial advances during the Pleistocene did not extend into Kentucky and Tennessee and this region likely served as a refuge for both terrestrial and aquatic fauna (Jacquemin 2016). As glaciers retreated, populations at the northern limits of these refugia would have advanced northward into newly habitable territory. These expansions would have left a genetic signature characterized by reduced genetic variation in the northern end of their distribution as a result of founder effects. Repeated expansions of *A. barbouri* northward from Tennessee may explain the polyphyletic relationships of contemporary Tennessee *A. barbouri* with respect to populations in the north. Specifically, mitochondrial *A. barbouri* populations in northeastern Kentucky, Ohio and Indiana (Clade V) are nested within Tennessee *A. barbouri* as are mitochondrial haplotypes from Ambystomatid unisexuals and *A. texanum*. Genetic evidence of repeated northward expansions has been reported for the Eastern Woodrat (*Neotoma floridana*; Hayes & Harrison 1992) and other southeastern fauna (see Hewitt 1995 for review).

### Management Implications and Conclusions

Results from our genomic survey have specific implications for the design of management and conservation strategies that may improve the long-term persistence of *A. barbouri* in Tennessee. First, patterns of genetic variation in mitochondrial and nuclear genomes support the assignment of three genetically distinct units for management that warrant the designation of evolutionary significant units (ESUs), where ESUs are defined as groups of populations that show phylogeographic differentiation for mtDNA haplotypes and divergence in allele frequencies in nuclear markers (Moritz 1994, 1999). These three units include a Northern ESU encompassing all Sumner County populations (S2, S5, S7 and S8), a Central ESU that includes populations from eastern Davidson and Wilson Counties (D3, W1, W3 and W4), and a Southern ESU that includes populations in Williamson, Rutherford, and Bedford Counties (Wil2, B6, R1, R7, and R9). Patterns of differentiation between these three groups of populations suggest a long history of genetic isolation at both mitochondrial and nuclear markers, such that different groups are likely to possess unique combinations of adaptive genetic variation and have likely experienced independent evolutionary trajectories.

Evolutionarily and phylogenetically distinctive populations contribute disproportionately to genetic diversity and should be ranked highly in regards to conservation priority. Geographically peripheral populations are frequently observed to be genetically less-redundant than more central populations and are more likely to possess potentially adaptive genetic variation (Volkmann et. al. 2014). For *A. barbouri,* Tennessee populations are geographically disjunct from the majority of *A. barbouri* breeding populations (Niemiller et al. 2006) and occupy the southernmost edge of the species distribution. Results from this study suggest that Tennessee *A. barbouri* should be prioritized for conservation planning as these populations are both genetically diverse and evolutionarily distinct from populations in the northern part of their distribution. The distinctiveness of these populations is further evidenced by observed differences in reproductive life-history traits, including mean diameter of early stage ovum size and number of eggs per clutch (Niemiller et al. 2009). Prioritizing peripheral populations with adaptive genetic variation and evolutionary potential is even more critical when considering environmental challenges that accompany climate change. Amphibians in general are very sensitive to climate change as their reproductive life histories are linked to temperature and precipitation (Corn 2005). However, there is evidence that populations at the warm-range edge of their distribution are more resilient (Razgour et al. 2019). Protection of *A. barbouri* breeding sites in Tennessee may be instrumental to ensuring the long-term viability of this species as a whole.

## Acknowledgments

We thank Tennessee Wildlife Resources Agency (TWRA) for providing funding to support this work. Ryan Hanscom assisted with sample collections. Joanna Bellan assisted with mitochondrial sequencing and data analysis. Andrea Drayer provided *A. barbouri* tissue samples from Kentucky populations. We also thank Kristin Womble for producing the map in Figure 2.

## References

Allio R, Donega S, Galtier N, Nabholz B. Large variation in the ratio of mitochondrial to nuclear mutation rate across animals: implications for genetic diversity and the use of mitochondrial DNA as a molecular marker. Mol. Biol. Evol. 2017; 34(11):2762–2772.

Anderson MA, Campbell JR, Carey AN, Dodge DR, Johnston RA, Mattison ER, Seddon RJ, Singer NL, Miller BT. Population survey of the streamside salamander in the Nashville Basin of Tennessee. Southeast Nat. 2017: 13(1):101–107.

Toews DP, Brelsford A. The biogeography of mitochondrial and nuclear discordance in animals. Molecular ecology. 2012: 21(16):3907–30.

Bogart JP, Bi K, Fu J, Noble DW, Niedzwiecki, J. Unisexual salamanders (genus *Ambystoma*) present a new reproductive mode for eukaryotes. Genome 2007; 50(2):119–136.

Catchen JM, Amores A, Hohenlohe P, Cresko W, Postlethwait JH. *Stacks*: building and genotyping loci *de novo* from short-read sequences. G3-Genes Genom Genet 2011; 1(3):171–182.

Charney ND, Ireland AT, Bettencourt BR. Mapping genotype distributions in the unisexual *Ambystoma* complex. J Herpetol 2014; 48(2):210–219.

Church SA, Kraus JM, Mitchell JC, Church DR, Taylor DR. Evidence for multiple Pleistocene refugia in the postglacial expansion of the eastern tiger salamander, *Ambystoma tigrinum*. Evolution 2003; 57(2):372–383.

Corn, P.S., 2005. Climate change and amphibians. Biodiv. Cons. 2005; 28(1).

Do C, Waples RS, Peel D, Macbeth GM, Tillett BJ, Ovenden JR. NeEstimator v2: re-implementation of software for the estimation of contemporary effective population size (Ne) from genetic data. Mol. Ecol. Resour. 2014; 14(1):209–214.

Eaton DAR. PyRAD: Assembly of de novo RADseq loci for phylogenetic analyses. Bioinformatics 2014; 30(13):1844–1849.

Earl DA, VonHoldt BM. Structure Harvester: a website and program for visualizing STRUCTURE output and implementing the Evanno method. Conserv. Genet. Resour. 2012; 4(2):359–361.

Eastman JM, Niedzwiecki JH, Nadler BP, Storfer A. Duration and consistency of historical selection are correlated with adaptive trait evolution in the streamside salamander, *Ambystoma barbouri*. Evolution 2009; 63(10):2636–2647.

Elshire RJ, Glaubitz JC, Sun Q, Poland JA, Kawamoto K, Buckler ES, Mitchell SE. A Robust, Simple Genotyping-by-Sequencing (GBS) Approach for High Diversity Species. PLoS One. 2011; 6(5):e19379.

Evanno G, Regnaut S, Goudet, J. Detecting the number of clusters of individuals using the software STRUCTURE: a simulation study. Mol. Ecol. 2005; 14:2611–2620.

Excoffier L, Smouse PE, Quattro JM. Analysis of molecular variance inferred from metric distances among DNA haplotypes: application to human mitochondrial DNA restriction data. Genetics 1992; 131(2):479–491.

Excoffier L, Lischer HEL. Arlequin suite ver 3.5: a new series of programs to perform population genetics analyses under Linux and Windows. Mol. Ecol. Resour. 2010; 10(3):564–567.

Funk WC, Tallmon DA, Allendorf FW. Small effective population size in the long-toed salamander. Mol Ecol 1999; 8(10):1633–1640.

Fusco, N.A., Pehek, E. and Munshi-South, J. Urbanization reduces gene flow but not genetic diversity of stream salamander populations in the New York City metropolitan area. Evol. App. 2010; 14(1): 99–116.

Geist J. Strategies for the conservation of endangered freshwater pearl mussels (*Margaritifera margaritifera L*.): a synthesis of conservation genetics and ecology. Hydrobiologia 2010; 644(1):69–88.

Haasl RJ, Payseur BA Multi-locus inference of population structure: a comparison between single nucleotide polymorphisms and microsatellites. Heredity 2011; 106(1):158–171.

Hall TA. BioEdit: a user-friendly biological sequence alignment editor and analysis program for Windows 95/98/NT. Nucl Acid 1999; 41:95–98.

Hare MP. Prospects for nuclear gene phylogeography. Trends Ecol. Evol. 2001; 16(12):700–706. https://doi.org/10.1016/S0169-5347(01)02326-6

Hartl DL, Clark AG. Principles of population genetics.1997; Sinauer Assoc. Inc, Sunderland, Massachusetts.

Hayes JP, Harrison RG. Variation in mitochondrial DNA and the biogeographic history of woodrats (Neotoma) of the eastern United States. Systematic Biology. 1992; 41(3):331–44.

Heredia-Bobadilla RL, Monroy-Vilchis O, Zarco-González MM, Martínez-Gómez D, Mendoza-Martínez GD, Sunny A (2016) Genetic structure and diversity in an isolated population of an endemic mole salamander (*Ambystoma rivulare* Taylor, 1940) of central Mexico. Genetica 2016; 144(6): 689–698.

Hewitt GM. Some genetic consequences of ice ages, and their role in divergence and speciation. Biological journal of the Linnean Society. 1996;58(3):247–76.

Hewitt G. The genetic legacy of the Quaternary ice ages. Nature. 2000; 405(6789):907–13.

Huang H, Knowles, LL. Unforeseen consequences of excluding missing data from next-generation sequences: simulation study of RAD sequences. Sys. Bio. 2016; 65(3): 357–365.

Huelsenbeck JP, Ronquist F. MRBAYES: Bayesian inference of phylogenetic trees. Bioinformatics 2001; 17(8): 754–755.

Jacquemin SJ, Ebersole JA, Dickinson WC, Ciampaglio CN. Late Pleistocene fishes of the Tennessee River Basin: an analysis of a late Pleistocene freshwater fish fauna from Bell Cave (site ACb-2) in Colbert County, Alabama, USA. PeerJ. 2016; 2(4):e1648.

Jehle R, Arntzen JW. Microsatellite markers in amphibian conservation genetics. Herpetol J 2002; 12:1–9.

Jombart T, Ahmed I. Adegenet 1.3-1: new tools for the analysis of genome-wide SNP data. Bioinformatics 2011; 27(21): 3070–3071.

Jombart T, Collins C. A tutorial for discriminant analysis of principal components (DAPC) using adegenet 2.0.0. London, Imperial College London, MRC Centre for Outbreak Analysis and Modelling. 2015.

Kardos M, Armstrong E, Fitzpatrick S, Hauser S, Hedrick P, Miller, J., Tallmon, D.A. and Funk, W.C., 2021. The crucial role of genome-wide genetic variation in conservation.

Keenan K, McGinnity P, Cross TF, Crozier WW, Prodöhl PA. diveRsity: An R package for the estimation of population genetics parameters and their associated errors, Methods in Ecology and Evolution. 2013; 4(8): 782–788.

Kopelman, Naama M., Jonathan Mayzel, Mattias Jakobsson, Noah A. Rosenberg, and Itay Mayrose. “Clumpak: a program for identifying clustering modes and packaging population structure inferences across K.” Molecular ecology resources 15, no. 5 (2015): 1179–1191.

Kraus F, Petranka JW. A new sibling species of *Ambystoma* from the Ohio River drainage. Copeia 1989; 94–110.

Micheletti SJ, Storfer A. An approach for identifying cryptic barriers to gene flow that limit species’ geographic ranges. Mol Ecol 2017; 26(2):490–504.

Miller MA, Pfeiffer W, Schwartz T. The CIPRES science gateway: a community resource for phylogenetic analyses. InProceedings of the 2011 TeraGrid Conference: extreme digital discovery 2011; Jul 18: 1–8.

Moritz C. Defining ‘evolutionarily significant units’ for conservation. Trends Ecol. Evol. 1994: 9(10):373–375.

Moritz, C. Conservation units and translocations: strategies for conserving evolutionary processes. Hereditas 1999; 130(3):217–228.

Niedzwiecki JH. Evolutionary history and hybridization of two mole salamander sister species from different habitats. Dissertation, University of Kentucky. 2005

Nichols R. Gene trees and species trees are not the same. Trends in Ecology & Evolution. 2001;16(7):358–64.

Niemiller ML, Glorioso BM, Nicholas C, Phillips J, Rader J, Reed E, Sykes KL, Todd J, Wyckoff GR, Young EL, Miller BT. Status and distribution of the Streamside Salamander, *Ambystoma barbouri*, in middle Tennessee. Am. Midl. Nat. 2006;156(2):394–399.

Niemiller ML, Glorioso BM, Nicholas C, Phillips J, Rader J, Reed E, Sykes KL, Todd J, Wyckoff GR, Young EL, Miller BT. Notes on the reproduction of the Streamside Salamander, *Ambystoma barbouri*, from Rutherford County, Tennessee. Southeast Nat. 2009; 8(1):37–44.

Nunziata SO, Lance SL, Scott DE, Lemmon EM, Weisrock DW. Genomic data detect corresponding signatures of population size change on an ecological time scale in two salamander species. Mol. Ecol. 2017 26(4):1060–1074.

Nunziata SO, Weisrock DW Estimation of contemporary effective population size and population declines using RAD sequence data. Heredity 2018; 120(3): 196–207.

Patton AH, Margres MJ, Epstein B, Eastman J, Harmon LJ, Storfer A. Hybridizing salamanders experience accelerated diversification. Scientific reports. 2020 Apr 16;10(1):1–2.

Posada D, Crandall KA. Modeltest: testing the model of DNA substitution. Bioinformatics (Oxford, England) 1998; 14(9):817–818.

Pritchard JK, Wen W, Falush D. Documentation for STRUCTURE software: Version 2. Department of Human Genetics, University of Chicago, Chicago. 2003

Razgour O, Forester B, Taggart JB, Bekaert M, Juste J, Ibáñez C, Puechmaille, SJ, Novella-Fernandez R, Alberdi A, Manel S. Considering adaptive genetic variation in climate change vulnerability assessment reduces species range loss projections. Proc Nat Acad Sci 2019.; 116(21):10418–10423.

Reis M and Yang Z. In: Anisimova M (ed.) Evolutionary Genomics. Methods in Molecular Biology, vol 1910. Humana, New York, NY. 2019

Savage WK, Fremier AK, Bradley Shaffer H. Landscape genetics of alpine Sierra Nevada salamanders reveal extreme population subdivision in space and time. Mol. Ecol. 2010 19(16):3301–3314.

Schmidt C, Garroway CJ. The population genetics of urban and rural amphibians in North America. Mol. Ecol. 2021

Scott AF, Miller BT, Brown M, Petranka JW. Geographic distribution: *Ambystoma barbouri*. Herpetol. Rev. 1997; 28:155.

Shaffer HB, McKnight ML. The polytypic species revisited: genetic differentiation and molecular phylogenetics of the tiger salamander *Ambystoma tigrinum* (Amphibia: Caudata) complex. Evolution 1996; 50(1):417–433.

Tennessee Wildlife Resources Agency Rules of Biodiversity. Rules and Regulations for in Need of Management, Threatened, and Endangered Species. Chapter 1660-01-32. 2018.

Thompson JD, Gibson TJ, Higgins DG. Multiple sequence alignment using ClustalW and ClustalX. Curr. Protoc. Bioinformatics 2003; (1):2–3.

Vences, M. and Wake, D.B., 2007. Speciation, species boundaries and phylogeography of amphibians. Amph. Biol.; 2007: 2613–671.

Volkmann L, Martyn I, Moulton V, Spillner A, Mooers AO (2014) Prioritizing populations for conservation using phylogenetic networks. PLoS One 2014; 9(2):e88945.

Waples RS, Do C (2008) LDNE: a program for estimating effective population size from data on linkage disequilibrium. Mol. Ecol Res. 2008; 8: 753–756.

Wang IJ, Johnson JR, Johnson BB, Shaffer HB (2011) Effective population size is strongly correlated with breeding pond size in the endangered California tiger salamander, *Ambystoma californiense*. Conserv. Genet. 2011; 12(4):911–920.

Withers DI, Condict K, McCoy R. A Guide to the Rare Animals of Tennessee. Division of Natural Areas, Tennessee Department of Environment and Conservation, Nashville, Tennessee. 2009

Yang Z. The BPP program for species tree estimation and species delimitation. Current Zoology. 2015; 61(5):854–65.

Zamudio KR, Savage WK. Historical isolation, range expansion, and secondary contact of two highly divergent mitochondrial lineages in spotted salamanders (*Ambystoma maculatum*). Evolution 2003; 57(7):1631–1652.

